# The cerebellum promotes sequential foraging strategies and contributes to the directional modulation of hippocampal place cells

**DOI:** 10.1101/2023.02.04.526971

**Authors:** Lu Zhang, Julien Fournier, Mehdi Fallahnezhad, Anne-Lise Paradis, Christelle Rochefort, Laure Rondi-Reig

**Author notes:** These authors contributed equally (Lu Zhang, Julien Fournier). These authors contributed equally (Christelle Rochefort, Laure Rondi-Reig). Current address: Coulter Department of Biomedical Engineering, Georgia Institute of Technology and Emory, Atlanta, GA, United States. Correspondence (LRR); (CR); (JF).

## Abstract

The cerebellum contributes to goal-directed navigation abilities and place coding in the hippocampus. Here we investigated its contribution to foraging strategies. We recorded hippocampal neurons in mice with impaired PKC-dependent cerebellar functions (L7-PKCI) and in their littermate controls while they performed a task where they were rewarded for visiting a subset of hidden locations. We found that L7-PKCI and control mice developed different foraging strategies: while control mice repeated spatial sequences to maximize their rewards, L7-PKCI mice persisted to use a random foraging strategy. Sequential foraging was associated with more place cells exhibiting theta-phase precession and theta rate modulation. Recording in the dark showed that PKC-dependent cerebellar functions controlled how self-motion cues contribute to the selectivity of place cells for both position and direction. Thus, the cerebellum contributes to the development of optimal sequential paths during foraging, possibly by controlling how self-motion and theta signals contribute to place cells coding.

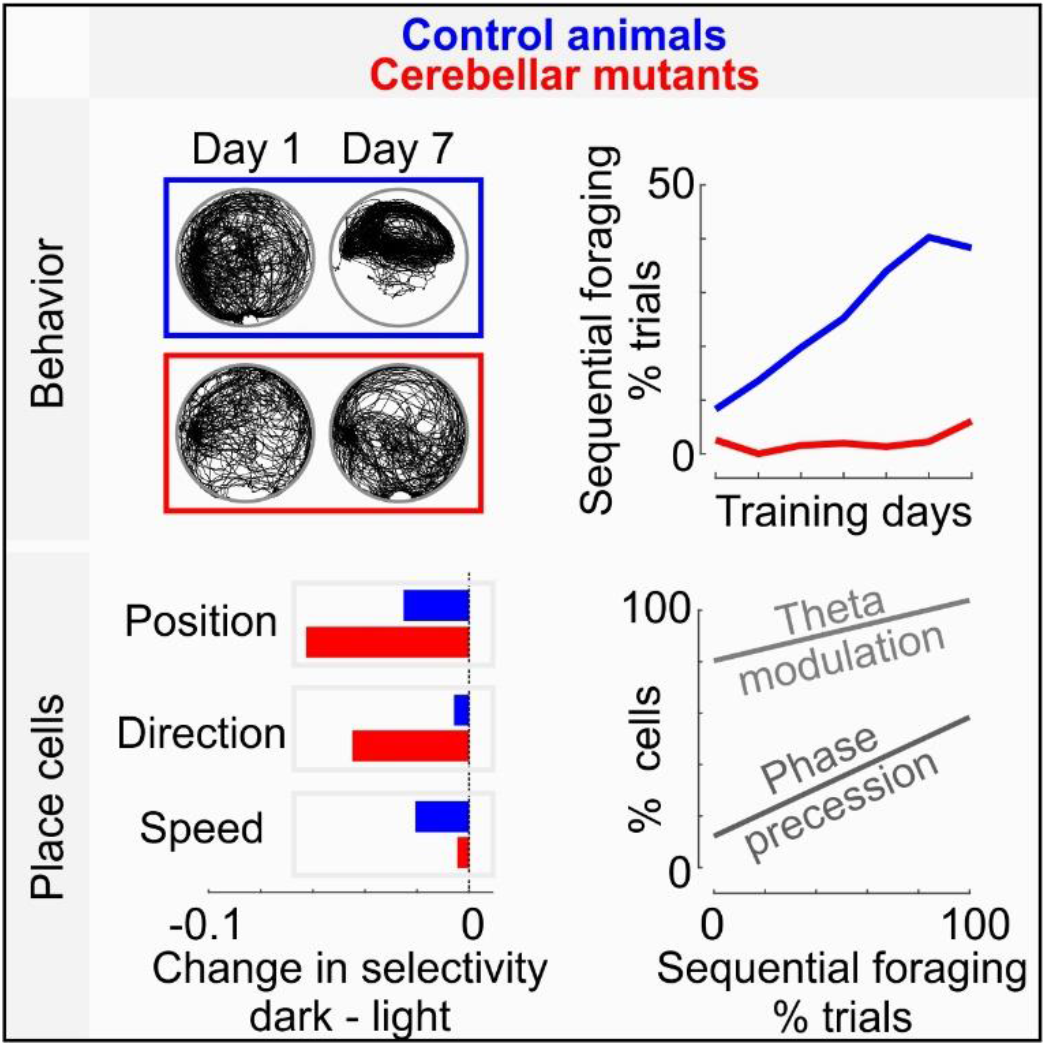

## Introduction

When foraging for food, animals optimize their foraging pattern to maximize the benefits while minimizing the energy cost^1^. This optimization is constrained by external factors such as the topology of the environment but also by physiological features of the animals that can limit their foraging efficiency. To spatially optimize their search trajectory, animals must estimate the distance and direction of possible targets from their current location and decide which target to go next. While the hippocampus is likely involved^2–4^, the nature of the hippocampal signals and the brain areas contributing to such spatial optimization of the foraging strategy remain elusive.

In the hippocampus, the position selectivity of place cells relies on the integration of sensory evidence from external landmarks and internal information related to the motion of the animal^5–8^. Previous research showed that the cerebellum is involved in how both types of signals contribute to hippocampal place fields: impairment of PKC-dependent cerebellar functions disrupt place fields when the animal relies on self-motion information^9^ while impairment of PP2B-dependent cerebellar functions results in a loss of stability of place fields relative to external cues^10^. Accordingly, mice with impaired PKC-dependent mechanisms exhibit learning deficits during goal-directed navigation^11^, particularly when they rely on self-motion information to learn a stereotyped trajectory between a fixed starting point and a goal location^9^.

Since the cerebellum and the hippocampus are co-activated during sequence-based navigation both in humans^12^ and mice^13^, we hypothesized that the cerebellum contributes to the development of spatially optimal foraging trajectories. To test this hypothesis, we designed a complex foraging task, where mice spontaneously developed different foraging strategies to obtain a reward after visiting a subset of hidden locations. We used L7-PKCI transgenic mice, which exhibit altered PKC-dependent mechanisms in cerebellar Purkinje cellsss^14,15^ and compared their behavior to that of littermate controls.

We found that the cerebellum promotes the spontaneous development of optimal sequential paths: during the first days of training, control mice quickly developed stereotyped spatial sequences to maximize their reward while L7-PKCI mice persisted to forage the environment randomly.

Hippocampal place cells were recorded in both groups after several weeks of training. By that time, some L7-PKCI mice had progressively developed a sequence-based strategy and various levels of optimization of the foraging behavior were represented across animals. This allowed us to investigate the features of hippocampal place cells associated with a sequence-based foraging strategy and over which cerebellum may act upon during early training days.

We found that the sequential foraging strategy was associated with more place cells exhibiting theta-phase precession and theta rate modulation. Moreover, a larger fraction of place cells tended to be modulated concomitantly by running direction and speed in animals using a sequence-based foraging strategy. Finally, by recording the same place cells in the dark, we found that cerebellum contributes to self-motion information underlying not just position but also direction signals encoded by the hippocampus. We conclude that the cerebellum promotes the development of optimal foraging strategies and that this contribution may reflect an impact of PKC-dependent cerebellar functions on self-motion and theta signals contributing to place cells coding.

## Results

To investigate the role of the cerebellum and the nature of the hippocampal signals associated with the optimization of foraging behaviors, we designed a complex foraging task, where mice were rewarded for visiting a subset of hidden locations. At each trial, two types of rewards had to be collected one after the other (Figure **1A**): the *foraging* reward was delivered upon visiting three different types of *foraging* sites (marked in *yellow*, *blue* and *purple* in Figure **1A**); the *goal* reward was then delivered when the animal returned to the *goal* zone (marked in *red* in Figure **1A**) and stayed there for more than 1s. None of the zones were visible to the animal. Rewards consisted of medial forebrain bundle stimulations so they were also invisible to the animal. Moreover, the foraging reward was delivered after a variable delay (0.6 to 2s) so that it was not associated with a precise spatial location. To complete the task, mice had to alternate between foraging and goal-directed behaviors but during the foraging phase, they were free to use any strategy to obtain the reward. In particular, they could either repeat the same trajectories, going through the same three zones in a sequence (Figure **1A**, *left*) or they could explore randomly the arena until they went through three different zones (Figure **1A**, *right*).

**Figure 1:**
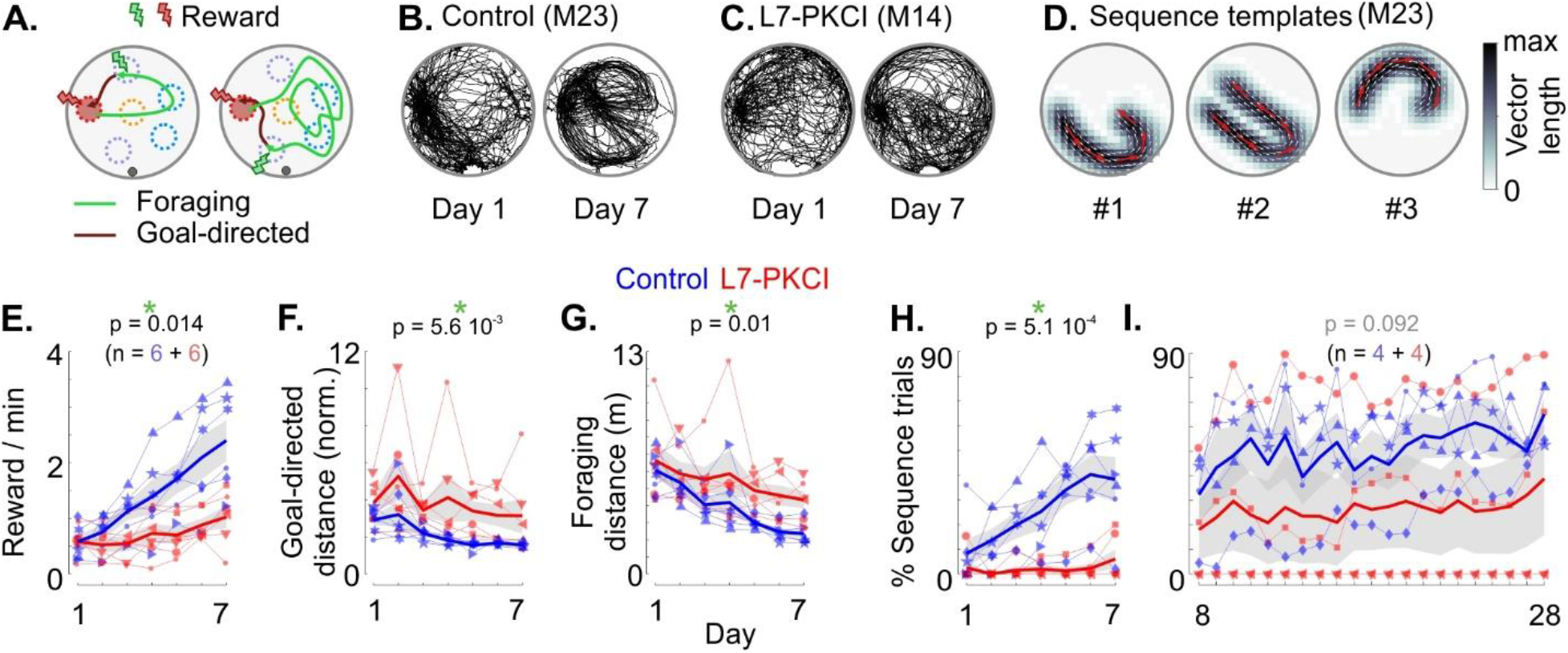
The cerebellum contributes to the spontaneous development of sequence-based foraging strategies. **A.** Schematic top view of the environment with successful trajectories. Mice were trained to collect two types of reward. The foraging reward (green) was delivered upon visiting all three types of foraging sites (each type being marked by a different color: yellow, purple and cyan); the goal reward (red) was then delivered when the animal returned and stayed in the goal zone (red) for 1s. None of the zones were visible to the animal. To collect the foraging reward, mice could repeat a spatial sequence, i.e. going systematically through the same three zones (left); or they could explore the arena randomly until they visited at least three different zones (right). **B.** Example of trials performed by a control mouse (M23) on the first (left) and seventh (right) day of training. **C.** Same as in B for a L7-PKCI mouse (M14). **D.** Trajectory templates identified by hierarchical clustering of trials from all sessions for one control mouse (M23). Arrows indicate mean direction of movement. Map intensity reflects the mean vector length in each position. Red arrows have been magnified (3x) to increase visibility. **E.** Number of rewards collected across the first 7 days of training for L7-PKCI (red, n = 6) and control (blue, n = 6) mice. Thick curves: mean performance across mice; gray area: +/− s.e.m. The p-value indicates the significance of the genotype effect (mixed GLM, LR test). **F.** Same as in E for the median distance traveled during the goal-directed trajectories. The goal-directed distance was normalized in each trial by the shortest distance between the position where the animal received the foraging reward and the position where it first entered the goal zone. **G.** Same as in E for the median distance traveled during the foraging trajectory. **H.** Same as E for the fraction of trials where the animal performed a spatial sequence (sequence trials). **I.** Fraction of trials where the animal performed a spatial sequence across later days of training (L7-PKCI, n = 4; Control, n = 4). Symbols correspond to different mice. See also Figure **S1**

### The cerebellum promotes the development of sequence-based foraging strategies

It was previously shown that mice with altered PKC-dependent cerebellar functions (L7-PKCI)^14,15^ exhibit deficits in using self-motion and external cues information in a goal-directed task^9,11^. However, it is unclear how PKC-dependent cerebellar functions contribute to the spatial optimization of the animal’s trajectory in a task where the reward can be obtained using different strategies. So we first asked whether L7-PKCI mice collected the foraging reward in our task using a strategy similar to control mice.

We trained six L7-PKCI transgenic mice and six littermate controls. Control mice rapidly learned the task: the number of collected rewards increased steadily during the first 7 days of training (Figure **1E**; mixed generalized linear model (GLM), likelihood ratio (LR) test of days effect: Δχ^2^(1) = 57.3, p = 3.6 10^−14^). This increase was associated with more direct trajectories back to the *goal* zone (Figure **1F**; mixed GLM, LR test of days effect: Δχ^2^(1) = 45.9, p = 1.2 10^−11^) and shorter traveled distances to collect the *foraging* reward (Figure **1G**; mixed GLM, LR test of days effect: Δχ^2^(1) = 62.0, p = 3.4 10^−15^). In contrast, the performances of L7-PKCI mice were lower (Figure **1E**; mixed GLM, LR test of genotype effect: Δχ^2^(1) = 5.99, p = 0.014), which could not be explained by differences in running speed (Figure **S1J** and **S1K**). They traveled a longer distance to collect the foraging reward (Figure **1G**, mixed GLM, LR test of genotype effect: Δχ^2^(1) = 6.64, p = 0.01). Their trajectories back to the goal zone were also less direct than those of control mice, consistent with previous reports^11^ (Figure **1F**, mixed GLM, LR test of genotype effect: Δχ^2^(1) = 7.68, 5.6 10^−3^).

During foraging, L7-PKCI and control mice developed different strategies (Figure **1B-1D** and **1H**). As performance increased, control mice performed more spatial sequences, i.e. they passed reliably over multiple trials through the same foraging zones to collect the *foraging* reward (Figure **1B**). We used a clustering approach to identify the trajectory templates that each animal was inclined to follow more frequently (Figure **1D** and **S1A**). We then identified for each training day, trials that matched one of those spatial sequences (*sequence trials*). In all control mice, our clustering approach identified one or more spatial sequence templates (Figure **S1C**) and the fraction of trials matching those sequences increased across training days (Figure **1H**, mixed GLM, LR test of days effect: Δχ^2^(1) = 160.1, p = 0). In contrast, L7-PKCI mice persisted to explore the arena more uniformly to collect the foraging reward (Figure **1C**, **1H** and **S1B**). While L7-PKCI mice showed a slight increase in performance associated with shorter foraging and goal-directed distances (Figure **1E**-**1G**), this improvement did not correlate with a development of sequence-based foraging (Figure **1C** and **1H**). We identified sequence trajectories in only two out of six L7-PKCI mice (Figure **S1D**) and the fraction of sequence trials performed by these mice remained much lower than in controls (Figure **1H**, mixed GLM, LR test of genotype effect: Δχ^2^(1) = 12.1, p = 5.1 10^−4^).

The sequential exploration performed by control mice likely corresponded to spatial sequences rather than stereotyped motor sequences. Some mice indeed alternated between multiple spatial sequences that were clearly different from one another, even within the same session (Figure **1B**, **1D** and **S1A-S1D**). Moreover, when animals missed one of the foraging zones in a sequence, they often repeated only some part of the sequence rather than started over the entire sequence from the starting point (Figure **S1E**).

The behavioral differences between L7-PKCI and control mice faded over prolonged training (Figure **1I** and **S1F-S1I**). Eight out of twelve mice (with functional tetrode implants; 4 L7-PKCI, 4 controls) were further trained beyond the first seven days period. During this prolonged training, 2/4 L7-PKCI mice progressively did more sequence trials, reaching levels similar to control mice (Figure **1I**, mixed GLM, LR test of genotype effect: Δχ^2^(1) = 2.85, p = 0.09). Their trajectories back to the goal zone also became as direct as in control mice (Figure **S1G**, mixed GLM, LR test of genotype effect: Δχ^2^(1) = 2.85, p = 0.092). Thus, while PKC-dependent cerebellar functions appear to promote the development of foraging sequences, other compensatory mechanisms may contribute to a delayed acquisition of such strategies in some L7-PKCI mice.

### Sequence-based foraging strategies correlate with theta modulation and theta-phase precession in the hippocampus

Hippocampal neurons were recorded after several weeks of training, when tetrodes had reached the hippocampal CA1 region. By that time, two out of four L7-PKCI mice had developed sequence-based strategies, similar to control mice (Figure **1I**). While this prevented us from finding a direct interaction between genotype, behavior and hippocampal coding, it allowed us to search for features of hippocampal place cells that were associated with various levels of optimization of the foraging strategy across animals (Figure **2A-2D**).

**Figure 2.**
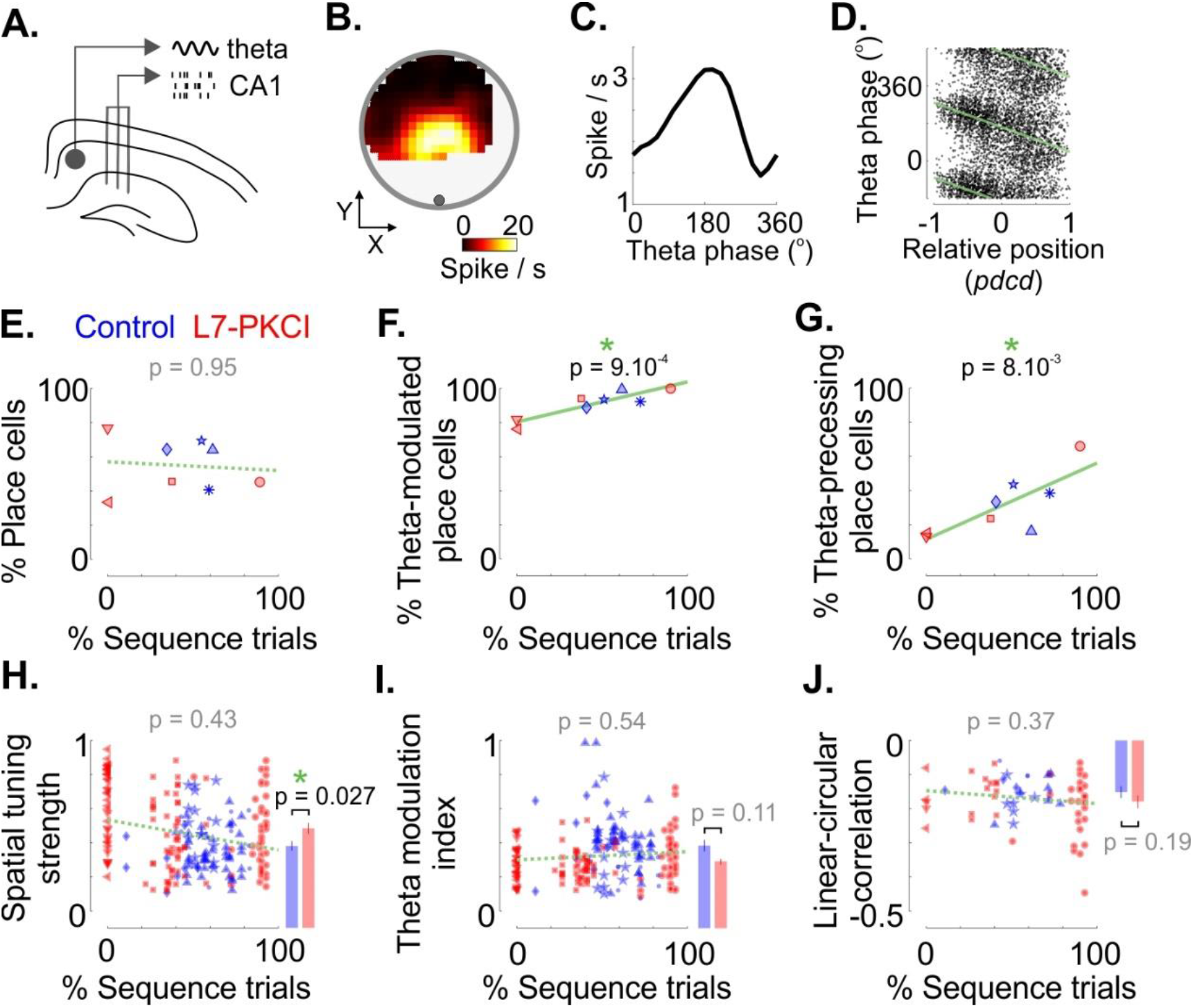
Sequence-based foraging behavior correlates with the modulation of hippocampal place cells by theta phases. **A.** Place cells were recorded with three tetrodes inserted in the CA1 region of the hippocampus. Theta oscillations were extracted from the local field potential recorded in the white matter above the hippocampus. **B.** Place field of an example cell from a mouse performing spatial sequences. **C.** Example of modulation of firing rate by the phase of the theta oscillation. **D.** Example of a theta-phase precessing place cell. Theta-phase precession was quantified by correlating the theta phase of the spikes and the relative position of the mouse within the neuron’s place field. The relative position was computed as the distance of the mouse to the center of mass of the place field, projected onto the current direction^21^ (*pdcd*). **E.** Percentage of place cells identified in L7-PKCI (*red*) and control (*blue*) mice as a function of the median percentage of sequence trials across training days. Total percentages: control, 59.7%; L7-PKCI, 45.0%. The p-value indicates the significance of the effect related to the fraction of sequence trials (mixed GLM, LR test). **F.** Fraction of place cells whose firing rate was significantly modulated across theta phases, expressed as a function of the median percentage of sequence trial for each animal. *Green line*: best fitting regression line. The p-value indicates the significance of the effect related to the fraction of sequence trials (mixed GLM, LR test). **G.** Same as in F for the fraction of place cells that exhibited significant theta-phase precession. **H.** *Left*, spatial tuning strength of L7-PKCI (*red*) and control (*blue*) place cells as a function of the percentage of sequence trials. The p-value indicates the significance of the effect related to the fraction of sequence trials (mixed GLM, LR test). *Right*, median tuning strength (+/− s.e.m.) across L7-PKCI (*red*) or control (*blue*) mice. The p-value indicates the significance of the genotype effect (mixed GLM, LR test). **I.** Same as in H for the strength of the modulation of firing rate by theta phase (*Theta modulation index*). **J.** Same as in H for the linear-circular correlation between the theta phase of the spikes and the position within the place field of each place cell. Symbols correspond to different mice. See also Figure **S2**.

Place cells were identified based on the spatial selectivity, stability and predictive power of their place field (135 in L7-PKCI, 80 in controls; out of 434 pyramidal neurons). The fraction of place cells was unrelated to the genotype or the foraging strategy of the animal (Figure **2E**, mixed GLM, LR test of the genotype effect: Δχ^2^(1) = 1.65, p = 0.20; LR test for the fraction of sequence trials: Δχ^2^(1) = 0.004, p = 0.95). At the single cell level, place cells appeared more selective in L7-PKCI mice than in controls (Figure **2H**, mixed GLM, LR test of the genotype effect: Δχ^2^(1) = 4.91, p = 0.027). However, their spatial selectivity was unrelated to whether animals performed more or less sequence trials (Figure **2H**, mixed GLM, LR test for the fraction of sequence trials: Δχ^2^(1) = 0.63, p = 0.43).

As expected, place cells were coupled to hippocampal theta oscillations: their firing rate oscillated at the scale of a theta cycle^16^ (Figure **2C**) and their spikes occurred at progressively earlier theta phases as the animal traversed their place field^17^ (theta-phase precession, Figure **2D**). Theta oscillations and theta-phase precession have previously been related to navigational performance and trajectory planning^18–20^. We thus asked whether the modulation of place cell firing by theta phase reflected the differences in foraging strategies observed across animals.

We found that the sequential foraging strategy was associated with more place cells coupled to the theta oscillation (Figure **2F-2G**). Indeed, the fraction of theta-modulated place cells was positively correlated with the fraction of sequence trajectories (Figure **2F**, mixed GLM, LR test for the fraction of sequence trials: Δχ^2^(1) = 11.09, p = 8.7 10^−4^) and the fraction of place cells exhibiting theta phase precession increased in animals performing a larger fraction of sequence trials (Figure **2G**, **S2A** and **S2B** mixed GLM, LR test for the fraction of sequence trials: Δχ^2^(1) = 6.95, p = 8.4 10^−3^).

It was previously reported that the theta-phase precession of hippocampal place cells appears stronger when animals run on a linear track than when they forage randomly in a 2D environment^22^. One possibility may thus be that theta-phase precession is more prominent in animals using a sequence-based strategy because their trajectories span a lower dimensional subspace than when foraging randomly. Our controls showed that this is unlikely to be the case. First, we found a similar correlation when selecting only traversals that were straight, fast and going through the center of the place fields (Figure **S2C**, mixed GLM, LR test for the fraction of sequence trials: Δχ^2^(1) = 7.99, p = 4.7 10^−3^). Second, the correlation between the theta phase of the spikes and positions within the place field did not strengthen with the fraction of sequence trials (Figure **2J**, *left*, mixed GLM, LR test for the fraction of sequence trials: Δχ^2^(1) = 0.81, p = 0.37), unlike what is observed when switching from a 2D environment to a linear track^22^. Thus, the larger fraction of place cells exhibiting theta phase precession in animals performing more sequence trials is most likely due to the behavioral strategy rather than to the dimensionality of the exploration path.

The larger fraction of theta-modulated place cells could not be explained by the strength of the LFP oscillations: no correlation was found between the fraction of sequence trials and the power of the theta oscillations in the LFP (Figure **S2D**, mixed GLM, LR test for the fraction of sequence trials: Δχ^2^(1) = 0.014, p = 0.91) or the strength of the theta modulation of firing rates (Figure **2I**, *left*, mixed GLM, LR test for the fraction of sequence trial: Δχ^2^(1) = 0.38, p = 0.54). Moreover, the fraction of theta-modulated interneurons did not exhibit the same trend as place cells (Figure **S2E**).

Considering genotypes separately, we found that the correlation between the sequential behavior and the percentage of theta-modulated or theta-phase precessing cells was significant for L7-PKCI mice (mixed GLM, LR test for the fraction of sequence trials: Figure **2F**, Δχ2(1) = 9.86, p = 1.7 10-3; Figure **2G**, Δχ2(1) = 10.07, p = 1.5 10^−3^) but not for control mice (mixed GLM, LR test for the fraction of sequence trials: Figure **2F**, Δχ2(1) = 0.57, p = 0.45; Figure **2G**, Δχ2(1) = 0.15, p = 0.69). Nonetheless, this may be due to the homogeneity of the behavior of control mice, for which the percentages of sequence trials span a range that may be too narrow to observe a significant correlation with the fraction of theta-modulated or theta-phase precessing cells.

Together, our analyses thus show that the development of a sequence-based foraging strategy is associated with more place cells exhibiting theta rate-modulation and theta-phase precession. Our results thus suggest that one way the cerebellum may promote sequence-based foraging strategies during early training days (Figure **1H**) is by contributing to the coupling between hippocampal place cells and theta oscillations.

### Sequence-based foraging strategies correlate with place cells modulation by direction and speed

Place cells are selective to spatial positions and theta phases but their activity is also modulated by movement direction and speed. It was previously reported that the directional modulation of place cells is contextual, as it may depend on the environment and on the task performed by the animal^23–26^. Since sequence-based foraging is characterized by trajectories where the direction of motion and spatial positions are related to each other in a reliable manner, we asked whether the modulation of place cells by the direction of movement reflected the ability of mice to perform reliable spatial sequences.

To estimate the selectivity of place cells to position (P), direction (D) and speed (S) while accounting for intrinsic correlations between those behavioral covariates, we used a Poisson Model (GLM, Figure **3A**) whereby each cell response is assumed to result from a multiplicative contribution of one or several covariates. Furthermore, to test for the significance of the contribution of each covariate, we used a forward selection procedure and identified the simplest model (i.e. with the fewest covariates) that resulted in a significant improvement of the prediction accuracy^27^.

**Figure 3.**
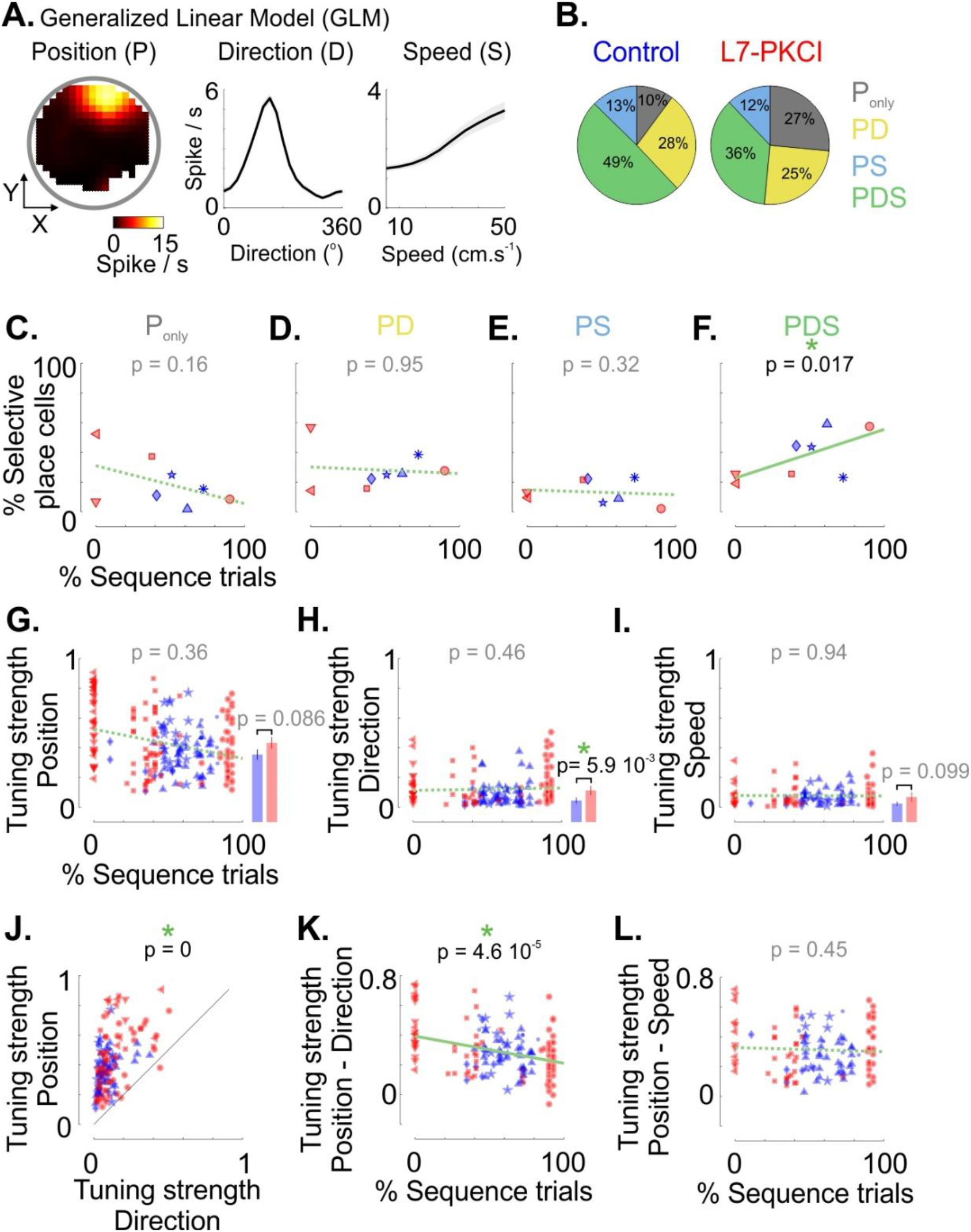
Sequence-based foraging behavior correlates with the modulation of hippocampal place cells by movement direction and speed. **A.** Tuning for position (P), movement direction (D) and running speed (S) estimated by a Poisson generalized linear model (GLM). **B.** Fractions of place cells in control (*left*) and L7-PKCI (*right*) mice that were identified as selective to position only (P_only_), position and movement direction (PD), position and running speed (PS) or position, movement direction and running speed (PDS). **C.** Percentage of place cells selective to position only as a function of the percentage of sequence trials (median across training days). The p-value indicates the significance of the effect related to the fraction of sequence trials (mixed GLM, LR test). **D.** Same as in C for place cells selective to position and movement direction. **E.** Same as in C for place cells selective to position and running speed. **F.** Same as in C for place cells selective to position, movement direction and speed. **G.** *Left*, strength of tuning to spatial positions for L7-PKCI (red) and control (*blue*) place cells, as a function of the percentage of sequence trials. The p-value indicates the significance of the effect related to the fraction of sequence trials (mixed GLM, LR test). *Right*, histograms report the median (+/− s.e.m.) across L7-PKCI (*red*) and control (*blue*) place cells. The p-value indicates the significance of the genotype effect (mixed GLM, LR test). **H.** Same as in G for the tuning to movement direction. **I.** Same as in G for the tuning to running speed. **J.** Tuning strength to position as a function of the tuning strength to movement direction. The p-value indicates the significance of the covariate effect, i.e. position vs. direction (mixed GLM, LR test). **K.** Difference between the tuning strength to position and direction as a function of the percentage of sequence trials. The p-value indicates the significance of the interaction between the covariate and the fraction of sequence trials (mixed GLM, LR test). **L.** Same as in K for the difference between tuning strength to position and speed. Symbols correspond to different mice. See also Figure **S3**.

A substantial fraction of place cells was significantly modulated by the direction of movement (Figure **3B**, *PD + PDS*: ~65%, 142/215; L7-PKCI, 62%; controls, 77%) or by running speed (Figure **3B**, *PS + PDS*: ~50%; 113/215; L7-PKCI, 48%; controls, 62%). The fraction of place cells exhibiting a contribution by movement direction (*PD*), speed (*PS*), both (*PDS*) or none of them (*P_only_*) were similar in L7-PKCI and control mice (Figure **3C**-**3F**, mixed GLM, LR test for the genotype effect: P_only_, Δχ^2^(1) = 1.27, p = 0.26; *PD*, Δχ^2^(1) = 0.08, p = 0.77; *PS,* Δχ^2^(1) = 0.14, p = 0.71; *PDS*, Δχ^2^(1) = 1.03, p = 0.31; *(PD + PDS)*, Δχ^2^(1) = 0.36, p = 0.55; *(PS + PDS)*, Δχ^2^(1) = 1.59, p = 0.21).

The proportions of place cells modulated by either direction (*PD*) or speed (*PS*) were unrelated to the foraging strategy (Figure **3D** and **3E**; mixed GLM, LR test for the fraction of sequence trials: *PD*, Δχ^2^(1) = 4 10^−3^, p = 0.95; *PS*, Δχ^2^(1) = 0.98, p = 0.32). However, we found that the sequence-based foraging strategy was associated with a larger fraction of place cells that were concomitantly modulated by both direction and speed (Figure **3F** and **S3A**, mixed GLM, LR test for the fraction of sequence trials: *PDS*, Δχ^2^(1) = 5.66, p = 0.017) while conversely, place cells selective to position only (*P_only_*) tended to decrease in mice performing more sequence-based trajectories (Figure **3C**; mixed GLM, LR test for the fraction of sequence trial: *P_only_*, Δχ^2^(1) = 1.95, p = 0.16).

At the single cell level, the selectivity to direction was stronger in L7-PKCI mice than in controls (Figure **3H**; mixed GLM, LR test for genotype effect: Δχ^2^(1) = 5.66, p = 5.9 10^−3^) but tuning to position, movement direction or speed appeared unrelated to the fraction of sequence trials (Figure **3G**-**3I** and **S3B**; mixed GLM, LR test for the fraction of sequence trials: *Position*, Δχ^2^(1) = 0.83; *Direction*, Δχ^2^(1) = 0.54, p = 0.46; *Speed*, Δχ^2^(1) = 5.7 10^−3^, p = 0.94). Yet, we found that the relative strengths of position and direction tunings correlated with the foraging strategy: while place cells were more selective to positions than to direction or speed (Figure **3J** and **S3C**), the gap in tuning strength between position and direction decreased when the fraction of sequence trials increased (Figure **3J** and **3K**; mixed GLM, LR test for the interaction between covariate (*P* or *D*) and the fraction of sequence trials: Δχ^2^(1) = 16.6, p = 4.6 10^−5^). In contrast, no such relationship was found when comparing selectivity to position and speed (Figure **3L**; mixed GLM, LR test for the interaction between covariate (*P* or *D*) and the fraction of sequence trials: Δχ^2^(1) = 0.58, p = 0.45). Thus, the selectivity of place cells to directions, relative to their selectivity to positions, tended to be stronger in animals using a sequence-based foraging strategy than in those foraging randomly.

Although recordings were performed on later days of training, these results may suggest that one possible way the cerebellum could promote the early development of sequence-based foraging strategies (Figure **1H**) is by influencing the balance between position, direction and speed contributions to place cells coding.

### The cerebellum contributes to self-motion signals underlying direction modulation in the hippocampus

It was previously shown that in the dark, the position selectivity of L7-PKCI place cells is disrupted, demonstrating that the cerebellum participates in self-motion signals underlying the selectivity of place cells to positions^9^. Since place cells are also sensitive to the direction and speed of motion^23–26^, we asked whether cerebellum also contributes to the self-motion components underlying direction and speed signals encoded in the hippocampus and whether this contribution correlates with the foraging strategy used by the animals.

After the initial training period (7 days), training days were split into light and dark sessions (2 light sessions + 2 dark sessions + 1 light session). Mice were still able to perform the task in the dark condition, although their performance and the proportion of spatial sequences decreased compared to the light sessions (Figure **S4A** and **S4B**).

To compare hippocampal selectivity between light and dark conditions, we considered place cells which were stable in the light sessions before and after the dark condition (n = 186; L7-PKCI: n = 113; controls: n = 73) and used a Poisson model including position, direction and speed as possible covariates (Figure **3A**).

In the dark, L7-PKCI place cells exhibited a lower selectivity not only to positions but also to the direction of movement (Figure **4**). As expected^9^, the spatial selectivity of CA1 place fields decreased in the dark, significantly more for L7-PKCI mice than for controls (Figure **4C** and **4D**, mixed GLM, LR test for the interaction between light/dark condition and genotype: Δχ^2^(1) = 5.79, p = 0.016). But remarkably, in the dark, the selectivity of place cells to direction also decreased more in L7-PKCI mice than in controls (Figure **4A-4D** and **S4C**, mixed GLM, LR test for the interaction between light/dark condition and genotype: Δχ^2^(1) = 7.11, p = 7.7 10^−3^). In control mice, the modulation by movement direction was barely affected by the light/dark transition across the place cell population (Figure **4D**, mixed GLM, LR test for the light/dark effect: Δχ^2^(1) = 1.01, p = 0.32). In contrast, in L7-PKCI mice, the directional modulation of place cells was markedly decreased when turning the light off (Figure **4D**, mixed GLM, LR test for the light/dark effect: Δχ^2^(1) = 20.12, p = 7.3 10-6). Thus, PKC-dependent cerebellar functions play a role in the directional modulation of hippocampal place cells by weighting the relative contributions of self-motion information and external cues. The higher selectivity of L7-PKCI place cells in the light condition (Figure **3H** and **S4D**), which is consistent with previous reports^9,28^, could possibly be explained by the relatively stronger contribution of external cues in those animals, which may reduce encoding errors related to multiplexing external and self-motion information in the light condition.

**Figure 4.**
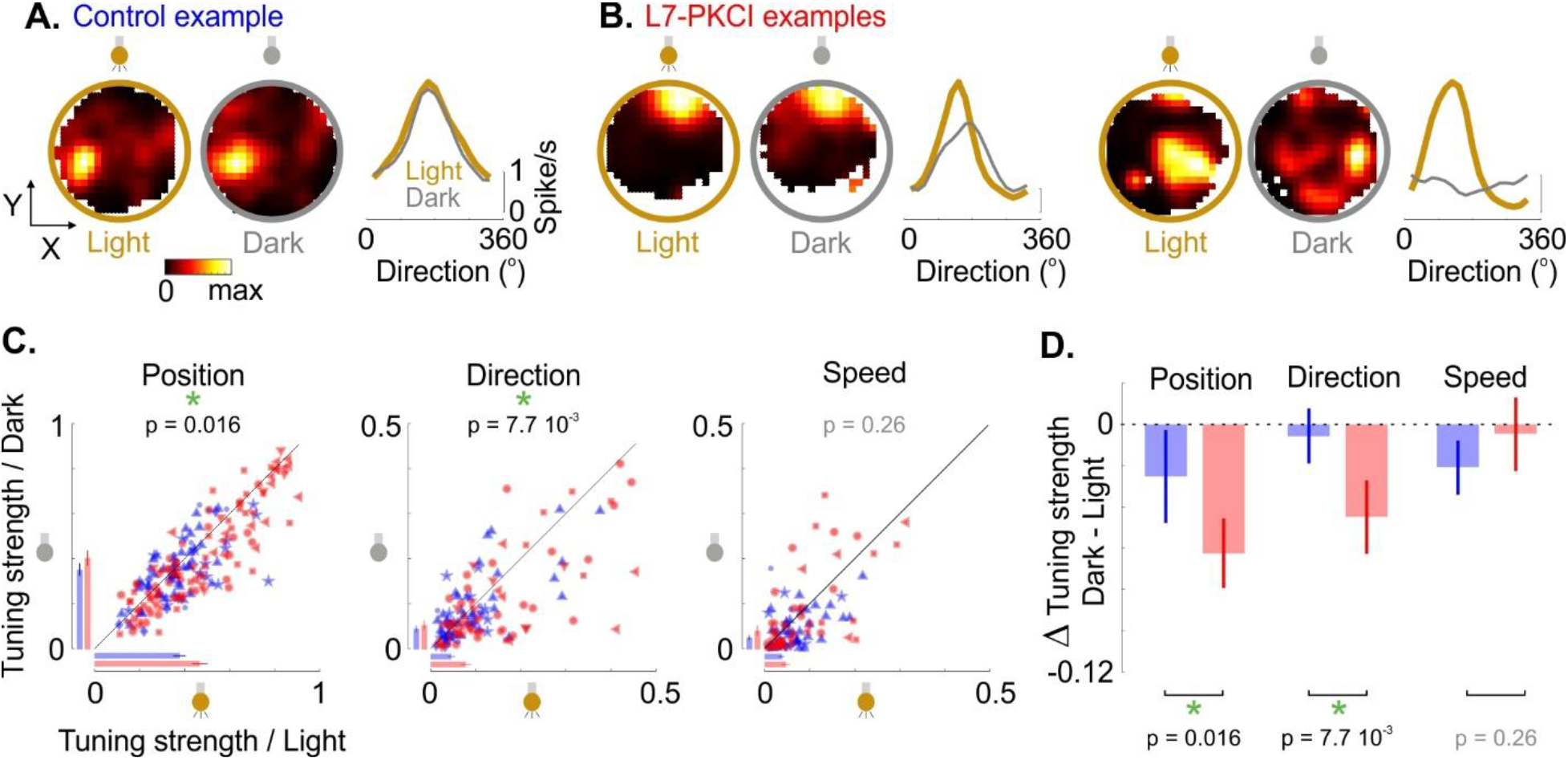
The cerebellum contributes to the directional modulation of place cells in the dark. **A.** Tuning to positions and direction during light (*gold*) and dark (*grey*) conditions for an example control place cell. **B.** Same as in A for two L7-PKCI place cell examples. **C.** Comparison of the tuning to position (*left*), movement direction (*middle*) or speed (*right*) between light and dark conditions, for L7-PKCI (*red*) and control (*blue*) place cells. Bars indicate the median across L7-PKCI (*red*) and control (*blue*) place cells (+/− s.e.m.). The p-values indicate the significance of the interaction between light/dark condition and genotype (mixed GLM, LR test). See also Figure **S4C**. Symbols correspond to different mice. **D**. Median (+/− s.e.m.) of the difference in tuning strength between dark and light conditions for position, direction and speed tuning of L7-PKCI (*red*) and control (*blue*) place cells. The p-values indicate the significance of the interaction between light/dark condition and genotype (mixed GLM, LR test). See also Figure **S4**.

No significant interaction was found between the foraging strategy and the transition from light to dark: the effect of the dark condition on position, direction or speed selectivity in L7-PKCI mice thus appeared unrelated to the fraction of sequence trials performed by the animals, at least during later days of training, when hippocampal recordings were performed (Figure **S4E**; mixed GLM, LR test for the interaction between light/dark condition and fraction of sequence trials: *Position*, Δχ^2^(1) = 0.04, p = 0.83; *Direction*, Δχ^2^(1) = 0.21, p = 0.65; *Speed*, Δχ^2^(1) = 1.17, p = 0.28). However, it may still be that the cerebellar control over hippocampal self-motion signals played a role during the early days of training and contributed to the earlier development of the sequence-based foraging strategy in control animals compared to L7-PKCI mice (Figure **1H**).

## Discussion

Optimizing foraging strategies likely recruits a large network of brain areas involved in navigation and decision making. We developed a complex navigation task, where mice were free to use different strategies to visit a set of hidden locations and obtain a reward. In this task, we found that cerebellum contributes to the optimization of the exploration strategy by promoting the development of spatial sequences during early training. We also found that in the hippocampus, sequence-based foraging strategies are associated with more place cells coupled to theta oscillations. Moreover, place cells were more often modulated by movement direction and speed in animals performing more spatial sequences. Finally, we showed that the cerebellum controls how self-motion signals contribute to the selectivity of place cells to not only position but also movement direction.

One possible interpretation of the behavioral differences observed in our task may be to consider that a sequence-based strategy corresponds to a series of sub-goal locations. In this context, the fact that L7-PKCI mice are impaired in developing such strategy is reminiscent of their deficits in learning simpler goal-directed navigation tasks^9,11,29^. Our results build on those previous reports by showing that L7-PKCI mice are impaired in spontaneously organizing sub-goal locations into an optimal spatial sequence.

The foraging strategy of L7-PKCI mice differed from control mice during early training but some L7-PKCI mice later developed spatial sequences over prolonged training. Therefore, by the time we recorded hippocampal place cells (i.e. after several weeks of training), L7-PKCI and control mice exhibited various levels of sequence-based behavior, going from no sequence to ~90% of trials matching a spatial sequence. This allowed us to identify neural correlates of the sequence-based foraging strategy to which the PKC-dependent cerebellar function may have contributed during early training.

Our results show that more place cells had their firing rate entrained by the theta oscillation when mice acquired a sequence-based foraging strategy rather than performed a random exploration. Remarkably, more place cells also exhibited theta phase precession in animals that developed a sequence-based foraging strategy. At the population level, theta-phase precession results in theta sequences whereby place cells fire sequentially on a scale of a theta cycle (~125 ms) in the same order as they are activated along the trajectory followed by the animal on a timescale of seconds. Theta sequences have been shown to reflect the current goals of the animal and may thus be involved in trajectory planning and decision making^19,20^. The inability of L7-PKCI mice to develop a sequence-based strategy as rapidly as control mice may thus reflect an impact of PKC-dependent cerebellar functions on the stabilization of hippocampal theta sequences.

Place cells are also modulated by the direction of travel of the animal. Previous research indicates that this modulation is contextual: on a linear track, the firing of place cells often depends on the direction of travel^5,6,24,30^ while in 2D environments, this directional modulation is weak when the animal is foraging randomly^24,31^ but stronger when the animal runs to a goal location^26,32^ or is trained to follow a stereotyped path^25^. We found that a larger fraction of place cells was modulated concomitantly by movement direction and running speed in animals using a sequence-based strategy. Moreover, the strength of tuning to direction relative to position tended to increase with the fraction of spatial sequences, consistent with a relatively stronger modulation of positional responses by running direction in animals following a sequential exploration path. The modulation of place cells by direction and speed may thus contribute to the development of a sequence-based foraging strategy. In L7-PKCI mice, turning the light off markedly decreased the modulation of place cells by movement direction compared to control animals, thus showing that the cerebellum controls how self-motion information contributes not only to positional^9^ but also directional signals in the hippocampus. The cerebellum may thus promote the development of sequence-based foraging strategies by enabling a more accurate encoding of positions and directions based on self-motion cues.

Place cells are also modulated by the direction of the head^33–35^ but it is yet unclear whether the movement and head direction signals that modulate place cells have the same origin. Our data show that in control mice, the selectivity of place cells to movement direction is similar in light and dark conditions, suggesting a strong contribution of self-motion to movement direction signals in the hippocampus. This is in contrast with the modulation of place cells by head direction, which strongly depends on visual cues^33^. Movement and head direction signals reaching the hippocampus may thus correspond to two distinct functional streams.

Overall, our analyses thus suggest that the role of the cerebellum in the optimization of the foraging strategy may be due to its implication in making proper prediction based on self-motion or to the development of hippocampal theta sequences. While L7-PKCI mice show no deficit in vestibulo-dependent motor tasks^9^, an altered integration of translational movements by cerebellar Purkinje cells^36^ may corrupt the estimation of self-motion components. One additional possibility may be that L7-PKCI mice have a lower motivational level compared to control mice, consistent with the influence of the cerebellum on the reward circuit and the recruitment of this projection during exploration^37^. Further experiments are needed to test the contribution of each of these mechanisms to the observed behavioral deficit in spatial optimization of the foraging strategy.

Several regions of the cerebellum (particularly Crus I and Lobule VI) have been shown to project to the hippocampus^38^. This anatomical link probably relies on multiple intermediate structures (subcortical and cortical) connecting the cerebellum to the hippocampus via several pathways. One such pathway may influence theta rhythms in the hippocampus via a disynaptic connection to the medial septum^39^. Another pathway may contribute to the encoding of self-motion and direction signals via the retrosplenial cortex^39^. The retrosplenial cortex (RSC) is indeed involved in head direction coding^40^, distance coding^41^ and receives disynaptic inputs from cerebellar dentate nucleus^42^. It may thus be one of the possible intermediates of the cerebellar control over self-motion signals reaching the hippocampus^43,44^. Accordingly, PKC-dependent cerebellar functions have recently been shown to also contribute to self-motion signals underlying head direction coding in RSC^28^.

How cerebellum may contribute to self-motion estimation? Self-motion signals used for navigation originate from the integration of multiple sensory cues (e.g. optic flow, head rotation, proprioception) and internal information related to motor commands. Cerebellum is well suited to combine those multiple streams of information: it receives external and internal information from multiple sources^29,45–47^ and exhibit nonlinear integration of multi-modal inputs at the level of granule cells that may underlie the detection of congruent information^46,47^. Purkinje cells have also been shown to process ego-centered direction signals from the vestibular system into a frame of reference anchored to the direction of gravity^48,49^. Thus, the cerebellum exhibits anatomical and computational features that are crucial in disambiguating self-from external motion. Similar to how the cerebellum may control motor commands in pre-motor areas, computations performed by the cerebellar network on self-motion cues may serve as a calibrating signal broadcasted to the navigational areas of the forebrain. Our results show that this cerebellar signal impacts position and direction coding in the hippocampus and contributes to the spatial optimization of the animal’s trajectory.

## Acknowledgments

This work was supported by the Fondation pour la Recherche Médicale DEQ20160334907-France, by the National Agency for Research ANR-17-CE37-0015-01 and ANR-18-CE16-0010-02 (LRR), by the Human Frontier Science program CDA 00058 - 2019 (JF) and by the Institut Universitaire de France (CR). This work also benefited from the support of the CNRS, INSERM and Sorbonne Université. We thank all members of the CEZAME team, especially Aurelie Watilliaux for helpful support in ethical procedures. We thank C.I. De Zeeuw for providing the L7-PKCI mice. We gratefully acknowledge the IBPS animal facility staff for their support.

## Author contributions

Conceptualization: CR and LRR; Methodology: CR and LRR; Experiment Setup: CR and LZ; Experimentation: LZ and CR; Software, Investigation, and Analysis: JF, LZ, and MF; Validation: JF, LZ, MF, ALP, CR, LRR; Visualization: JF; Writing First Draft: JF; Revision, Review, and Editing: JF, LZ, MF, ALP, CR, LRR, Funding Coordination, Project administration, reagents and resources: LRR.

## Methods

### Experimental model

All experiments were performed in accordance with the European ethical committee (European directive 86-609) and approved by the French Committee of Ethics Charles Darwin N° 5 (project 896.01). Data were collected from 12 adult male mice: 6 L7PKCI mice and 6 wild-type littermate controls. All animals were bred in a C57BL/6 mouse strain background. In L7PKCI mice, the pseudo-substrate PKC inhibitor (PKCI) was expressed under the control of the pcp-2 (L7) gene promoter, ensuring a selective suppression of PKC-dependent mechanisms in Purkinje cells. Hippocampal functions (synaptic transmission and plasticity), known to be essential for navigation tasks, are normal in L7-PKCI mice^11^. Their behavior during vestibulo-dependent motor tasks is also normal, both in the light and in the dark^9^. Mice were housed in standard conditions (12h light/dark cycle with water and food ad libitum). Experiments were carried out in blind conditions with respect to genotypes.

### Surgical procedure

Mice were implanted under deep anesthesia (Ketamine, 10 mg/kg; Xylazine, 10 mg/kg) at 3-7 months of age with three movable tetrodes (4 twisted 25-μm nichrome wires) targeting the CA1 region of the right hippocampus (AP: −2.2 mm; ML: 2 mm), a reference wire in the white matter above CA1 (DV: −0.9mm) (50-μm tungsten wire fixed on the tetrode cannula) and one stimulating electrode (100-μm-diameter stainless steel) targeting the left medial forebrain bundle (MFB) (AP: −1.4 mm; ML, +1.2 mm; DV: −4.8 mm).

### Behavioral training

Mice were tested on a spatial navigation task in a circular arena (100-cm diameter, 40-cm high) surrounded by a black circular curtain. No external landmarks were present, except one single proximal cue attached to the wall of the arena (a 210 × 297mm blue card with a non-transparent bottle in front). Tracking of mice positions was done online at 25Hz using the Smart system (Panlab, Barcelona, Spain) which also controlled the reward delivery. Rewards consisted of stimulations of the medial forebrain bundle (MFB).

The task consisted in collecting two types of reward one after the other: a *foraging* reward and a *goal* reward. To obtain the *foraging* reward, mice had to visit three different types of zones in any order: a central one (Figure **1A**, *yellow*) and two peripheral ones (Figure **1A**, *blue* and *purple*). The *foraging* reward was delivered upon entry in the last visited zone, with a random delay of 0.6 to 2 s. The *goal* reward could then be collected after the *foraging* reward, after staying 1 s inside the goal zone (Figure **1A**, *red*). None of the rewarded zones were visible to the animal. Each type of reward could only be collected one after the other, so mice had to alternate between exploring the arena and returning to the goal zone to obtain their rewards.

All mice were tested in this task for 7 days, with 5 daily sessions with light on (S1-S5, 12 minutes each). 8 mice (4 controls, 4 L7PKCI) with functional tetrode implants were kept for further training and recordings. For these animals, we introduced sessions with light off on the 8^th^ day: 2 out of 5 sessions (S3-S4) were performed in the dark with the cue removed, while the other 3 sessions remained unchanged.

Before mice were tested in this spatial task, we pre-trained them to associate reward delivery with the exploration of an invisible zone. This pre-training consisted of 5 daily sessions (15 minutes each, interleaved with 5 minutes in the home cage) and was done in two steps. First, we trained them to collect a reward upon visiting a single zone and staying outside this zone for more than 5s. Once they collected more than 15 rewards in one session, we trained them to perform the same task (albeit with a different zone) and to stay 1s in the goal zone before getting the reward. Mice that collected more than 15 rewards in one session of this second pre-training phase were considered as pre-trained and started being tested on the full task described above on the next day.

Calibration of the intensity of the MFB stimulation was done before the pre-training phase, 5 days after implantation. To do so, mice were tested in a nose-poke task using a non-transparent box (25 × 10 × 30 cm) with two nose poke places (1.5-cm diameter holes) on the same wall. Visits to the left nose poke place were detected by a photodetector and triggered the MFB stimulation. The preference for the left versus right nose poke location was calculated as the fraction of time spent at the rewarded nose poke in a 100-s period after the first visit^50^. Decreasing voltages (from 6V to 0V, 0.25V steps) were tested to find a tradeoff between low voltage stimulation and mice performance. The optimal voltage was defined as the lowest intensity that resulted in an 80% preference for the rewarded nose poke place across 3 successive sessions, interleaved with 5 minutes resting periods in the home cage. This optimal intensity ranged from 2V to 4V depending on individual mice.

### Behavioral analysis

Trials were first defined from one *goal* reward to the next. For each trial, trajectories were then split into a *foraging* trajectory and a *goal-directed* trajectory. The *foraging* trajectory was defined from the delivery of the *goal* reward to the entry in the last zone before the *foraging* reward was delivered; the *goal-directed* trajectory was defined from the delivery of the *foraging* reward to the point where it entered the goal zone before getting the *goal* reward.

To quantify the performance of mice in the spatial task, we calculated several metrics: the number of rewards per minute, the foraging distance, the goal-directed distance and the fraction of sequence trials. These measurements were made both for sessions in light and dark conditions.

The foraging distance was measured as the median across trials of the distance traveled on the *foraging* trajectories. Similarly, the goal-directed distance was computed as the median of the distance traveled during the goal-directed trajectories, after normalization by the Euclidian distance between the start and end points of each trajectory.

To quantify the fraction of sequence trials, we developed a clustering method to identify templates of trajectory that the animal would do preferentially during foraging. To do so, we binned spatial positions in 5-cm bins and the direction of movement in 20° bins. For time points where the animal was in the same spatial bin over consecutive times (< 200 ms), directions were averaged together and transformed into a unit vector. For each trial, direction vectors falling into the same spatial bins were averaged together, resulting in a spatial map *M* of direction vectors that we expressed in the complex space.

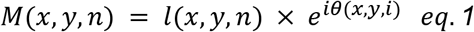

where *l* is the map of vector length across positions for trial *n* and *θ* is the map of directions.

Trials from all training days were clustered for each mouse using a hierarchical clustering approach to identify templates of reliable *foraging* trajectory. The distance *d*(*i*, *j*) between trials *i* and *j* was computed using the correlation between pairs of trials:

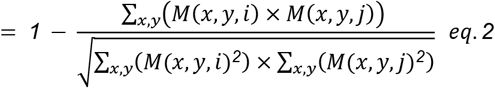

This distance matrix was used to build a dendrogram which was cut to identify clusters of trajectories that were more similar to one another. Clusters were considered as valid when they included more than 30 trials (over all training days). The cut-off threshold was adjusted between 1 – 0.6 and 1 – 0.9 to maximize the final number of valid clusters. Once these clusters were identified (between 0 and 3 depending on mice), sequence templates were computed as the average of all spatial maps of directions within each cluster. The *foraging* trajectory of a trial was then considered to match a specific sequence template when its Pearson’s correlation was higher than 0.6. The fraction of sequence trajectory was calculated for each training day as the number of trials matching a sequence template, divided by the total number of trials. For trials performed in the dark condition, we used the same templates as identified in the light condition.

### Electrophysiology

Extracellular signals from all electrodes were filtered through a unity-gain pre-amplifier (HS-16; Neuralynx, Bozeman, US). Spiking activity was sampled at 20kHz after filtering between 600 Hz and 9kHz. Local field potential (LFP) was filtered between 1 and 475 Hz and sampled at 1kHz. Data were digitized by a Power 1401 acquisition system controlled by Spike2 (CED, Cambridge, UK). MFB stimulation was delivered through a DS3 current stimulator (Digitimer, UK). Each MFB stimulation consisted of a 125 Hz train of 28 negative square wave pulses of 1-ms duration^51^.

5 days after implantation, tetrodes were slowly lowered in the brain (~60 μm per day) until reaching the hippocampus. Hippocampal recording was identified by increased multi-unit activity and sharp-wave ripples during resting periods.

Spikes were detected and sorted offline using Kilosort2^52^. Single units were considered as putative pyramidal neurons when they had a mean firing rate < 5Hz, a burst index > 2 and a prominent peak in their spike auto-correlogram between 3 and 18ms. Only well-isolated single units (isolation distance > 10) were considered for analysis.

For all single cell analyses, spikes and behavioral data were re-sampled at 100Hz and we only considered time points where the animals run speed was > 2.5 cm.s^−1^.

### Classical Place field

Place fields were estimated on light and dark sessions separately, by pooling the first two light sessions and the consecutive two dark sessions respectively.

Spatial positions were binned in 5-cm bins. Time points corresponding to position bins that were visited on less than 3 trials were excluded from the analysis. Spike count and occupancy were computed as a function of positions and both maps were smoothed with a Gaussian window (5-cm standard deviation). The classical place field of each neuron was then defined by dividing the spike count map with the occupancy map.

The spatial tuning strength was defined by measuring the sparsity of a place field^26,53^:

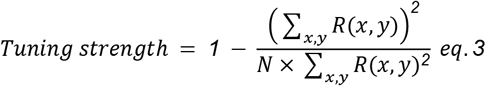

where *R*(*x*, *y*) is the mean firing rate estimated in position (*x*, *y*), and *N* is the number of bins.

To estimate the significance of spatial tuning strength, we measured place fields after circularly shifting spikes by a random delay (> 5s) for 500 times. Single units were considered as significantly tuned to positions when their spatial tuning strength was higher than the 95-percentile of the distribution of tuning strengths measured from the shuffle controls.

Predicted firing rates were computed using a 10-fold cross-validation procedure: 10 test sets were defined by splitting the data into non-overlapping subsets containing 10% of the data. The response of a neuron was predicted for each test set from the place field estimated with the remaining 90% of the data. To estimate the significance of the predictive power of a place field, we performed a likelihood ratio test comparing the place field prediction to the prediction by a constant mean firing rate. Neurons were considered to be significantly influenced by spatial position when the p-value of this test was < .05.

The stability of the place field over time was measured as the Pearson’s correlation between place fields estimated from different light or dark sessions, considering only positions where responses could be estimated in both sessions (i.e. with visits on more than 3 trials, see above).

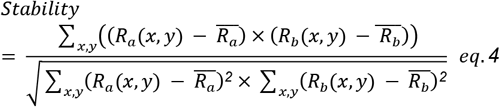

where *R*_*a*_ and *R*_*b*_ are the place fields estimated from sessions *a* and *b* and 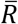 denotes the mean across positions.

A place field was considered as significantly stable when the correlation between place fields from the first and second light sessions (S1 and S2) was higher than the 95-percentile of correlations measured between place fields estimated in the same sessions from the shuffle controls. When comparing light and dark conditions, we considered neurons that were also significantly stable between the second and third light sessions (S2 and S5), to make sure that measured differences between light and dark conditions were not due to a progressive loss of stability of the neural response or recording.

Putative pyramidal neurons were identified as place cells when their place field met significance criteria for both tuning strength, predictive power and stability.

### Theta modulation and theta phase precession

Hippocampal theta oscillations were measured from the reference electrode placed in the white matter above the hippocampus. The broadband LFP signal was filtered between 6 Hz and 9 Hz. Peaks in the filtered signal were identified; those occurring less than 60 ms from the previous ones were discarded. Theta phases were measured as the fraction of time between successive peaks, converted into phases.

Firing rate modulation by theta oscillations was quantified by computing the tuning curve to theta phases. Theta phases were binned into 20° bins. Spike count and occupancy were measured as a function of theta phases and circularly smoothed by a Gaussian (20° standard deviation). The tuning of a neuron to theta phases was defined as the spike count vector divided by the occupancy vector.

The amplitude and peak phase of this theta modulation was measured by fitting a sinusoid to the estimated tuning curve. The theta modulation index was measured from the fitted sinusoid as:

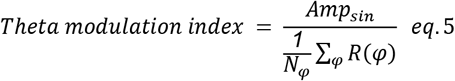

where *Amp*_*sin*_ denotes the amplitude of the best fitting sinusoid and *R*(*φ*) is the tuning curve across the *N*_*φ*_ phase bins.

To test for significance, we estimated similar tuning curves after circularly shifting spikes for 500 times by a random delay (> 5s). Neurons were considered as significantly modulated by theta phases when their theta modulation index was higher than the 95-percentile of the theta modulation indices measured from the shuffle controls.

To estimate theta phase precession from trajectories in a 2D environment, we measured the correlation between the theta phase of the spikes and the position of the animal within a place field^21^. In detail, positions falling within a place field were identified as those with a firing rate that was higher than the 95-percentile of firing rates estimated in the same positions from the shuffle control place fields (see above). Subfields were identified as areas with no common neighboring positions; when multiple subfields were identified, theta phase precession was measured only for the subfield with the largest area. Place field boundaries were defined as the convex hull including all significant positions falling in this subfield. For all spikes fired within these boundaries, we measured the distance of the animal to the center of mass of the field, projected on the run direction of the animal^21^. We then computed the correlation between this distance and the theta phase of spikes^54^ and only considered cells for further analysis when they fired at least 30 spikes within their field.

To test for significance of the correlation, we used the following statistics^54,55^:

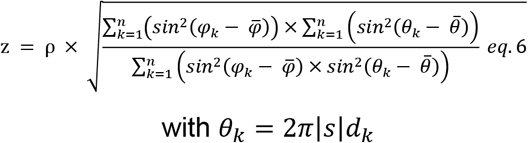

where ρ is the measured correlation, *s* is the estimated slope, *n* is the number of spikes, *φ*_*k*_ is the theta phase of spike *k* and *d*_*k*_ is the distance of the animal to the center of mass of the field projected on the run direction of the animal^21^. Since z is normally distributed for *n* sufficiently large, the p-value (*p*) can be calculated from the cumulative normal distribution:

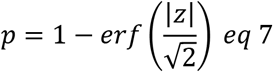

Neurons were considered to exhibit a significant theta phase precession when the theta phase × distance correlation had a p-value < .025 and when the theta phases covered across a traversal of the place field exceeded 45° ^21^.

To make sure that the observed differences in theta-phase precession were not just due to the higher reliability of place field traversals in animals using a sequence-based strategy (Figure **S2C**), we replicated the same analysis as described above after selecting only runs that were fast (i.e. minimal speed > 5 cm.s^−1^ and mean speed > 10 cm.s^−1^), straight (i.e. circular s.t.d. < 30°) and going through the center of the place field (i.e. −0.2 < pdcd < 0.2)^22^.

### Generalized Linear Model

To estimate the selectivity of place cells to multiple variables, we used a generalized linear model approach with a log link function^27^, assuming a multiplicative tuning to multiple covariates. Indeed, the spiking response of each neuron was expressed as the exponential of a weighted sum of one or more behavioral variables which could include position (P), direction of movement (D) or run speed (S) information:

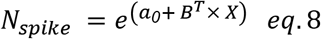

Where *N*_*spike*_ is a *N*_*time*_ × *1* vector of the number of spikes fired at every time *t*, *a*_*0*_ is a constant, *B* is a *N*_*time*_ × *N*_*B*_ matrix representing the behavioral state of the animal at time *t* with respect to the *N*_*B*_ variables included in the model (possibly including P, D or S) and *X* is a *N*_*B*_ × *1* vector describing the selectivity of the neuron to each of the variables included in the model. To construct *B*, spatial positions were binned into 5-cm bins; directions, into 20° bins and run speed, into 5 cm.s^−1^ bins. Columns in *B* corresponded to a succession of 0 and 1 describing in which of those bins the animal was at any time t. Time points corresponding to position bins that were visited on less than 3 trials were excluded from the analysis.

The best maximum likelihood solution (MLE) was estimated in Matlab, using the glmnet toolbox^56^ (with a final regularization parameter set to zero). The estimated coefficients (*X* in eq. 8) corresponding to each behavioral variable were smoothed (after exponentiation) with a Gaussian (5-cm s.d. for position; 20° for movement direction and 5-cm.s^−1^ for speed). The predictive power of the model was quantified using a 10-fold cross-validation procedure to measure the log-likelihood of held-out data under the model.

To estimate which variables (position, movement direction or speed) significantly contributed to the neuron’s response and thus had to be included in the model, we used a forward model selection procedure^27^. We first considered models with a single variable (P, H or S) and selected the one that resulted in the best predictive power (i.e. with the highest log-likelihood). We then added another variable to this model and identified the best two-variable model that resulted in a significant improvement of the prediction accuracy (compared to the best single variable model; likelihood ratio test, p < 0.05). We finally estimated the model with all three variables and tested if it resulted in a significant improvement of the prediction accuracy when compared to the best model identified at previous stages (likelihood ratio test, p < 0.05). The final optimal model was validated if it resulted in a significant improvement of the prediction accuracy over a constant mean model (likelihood ratio test, p < 0.05).

The estimated coefficients of the best model were converted into firing rate tuning curves^27^:

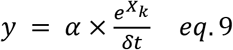

where 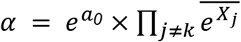 and *k* and *j* correspond to the covariates (P, D or S).

The tuning strength to each covariate was measured as:

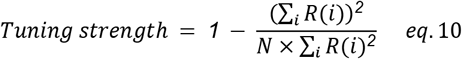

where *R* corresponds to the tuning to the considered variable across all bins.

### Statistical analyses

Statistical tests were performed by fitting generalized mixed effect models with nested random effects to account for variability across mice or sessions within mice (using Matlab function fitglme)^57^. The model was fitted using a Laplace approximation method and significance was estimated by means of likelihood ratio tests between nested models. The use of mixed models was justified by the fact that in all tests performed, including random effects always significantly improved the goodness of fit (LR test of random effects, p < 10^−10^). Data points further away from the median by 10 times the median absolute deviation were considered as outliers (in the limit of 5% of the data) and removed from statistical analysis.

To test for significant differences in behavioral performance across genotypes, we considered *genotype* as a fixed effect and *animals* and training *days* as nested random effect (i.e. *data ~ 1 + genotype + 1 | animal + 1 | animal:days*). This allowed us to test differences in behavioral performance between mouse lines while accounting for the variability across mice and training days. To test for the effect of training days on the performance of single mouse lines, we used *days* as a fixed effect and *animal* as random effect (i.e. *data ~ 1 + days + 1 | animal*).

For statistical analyses involving neural data in light condition, we considered the fraction of sequence trials (*sequence*) and *genotype* as fixed effects and animals as random effect (e.g. *data* ~ *1* + *sequence* + *genotype* + *1* | *animal*). When comparing results between light and dark sessions, we used light/dark condition (*L/D*), *genotype*, the fraction of sequence trials (*sequence*) and their cross-interactions (*L/D* × *genotype* and *L/D* × *sequence*) as fixed effects; *animals* and *neurons* were then included as nested random effects (*data* ~ 1 + *contrast* + *sequence* + *genotype* + *contrast:sequence* + *contrast:genotype +* 1 | *animal* + 1 | *animal:neuron*).

For all analysis involving proportions, we used a binomial distribution with a logit link function for the mixed model.

For all other analyses, we used a gamma distribution with a log link function as they all involved positive values with skewed distribution. Tuning strength data were mapped from the interval [0 1] to [0+inf[before fitting the model:

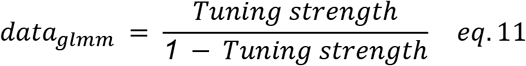

P-values were estimated using a likelihood ratio test (LR test) that compared the log likelihood of models with or without the tested variable. LR tests were performed within a forward search procedure. We first fitted all models including random effects and a single fixed variable (e.g. *contrast* or *genotype* or *sequence*). We then considered the fixed variable that led to the largest likelihood and computed its p-value with a LR test, relative to the model with no fixed effects. If the p-value was not significant (p > 0.05), we computed the p-values for the other fixed variables, using the model with no fixed effects as a reference (LR test). Otherwise, we included another fixed variable to the best single-variable model; selected the two-variable model that led to the largest likelihood and tested the significance of adding the second variable relative to the best single-variable model (LR test). If this p-value was not significant (p > 0.05), we computed the p-values for the remaining fixed variables, using the best single-variable model as a reference (LR test). Otherwise, we proceeded further by adding another fixed variable to the best two-variable model. We finally identified the best model (i.e. the one containing only variables that were identified as significant) and tested the significance of the interaction terms using a similar procedure. The significance of the interaction terms was always estimated after including the corresponding single variables to the equation of the model of reference.

## Supplementary Figures

**Figure S1:**
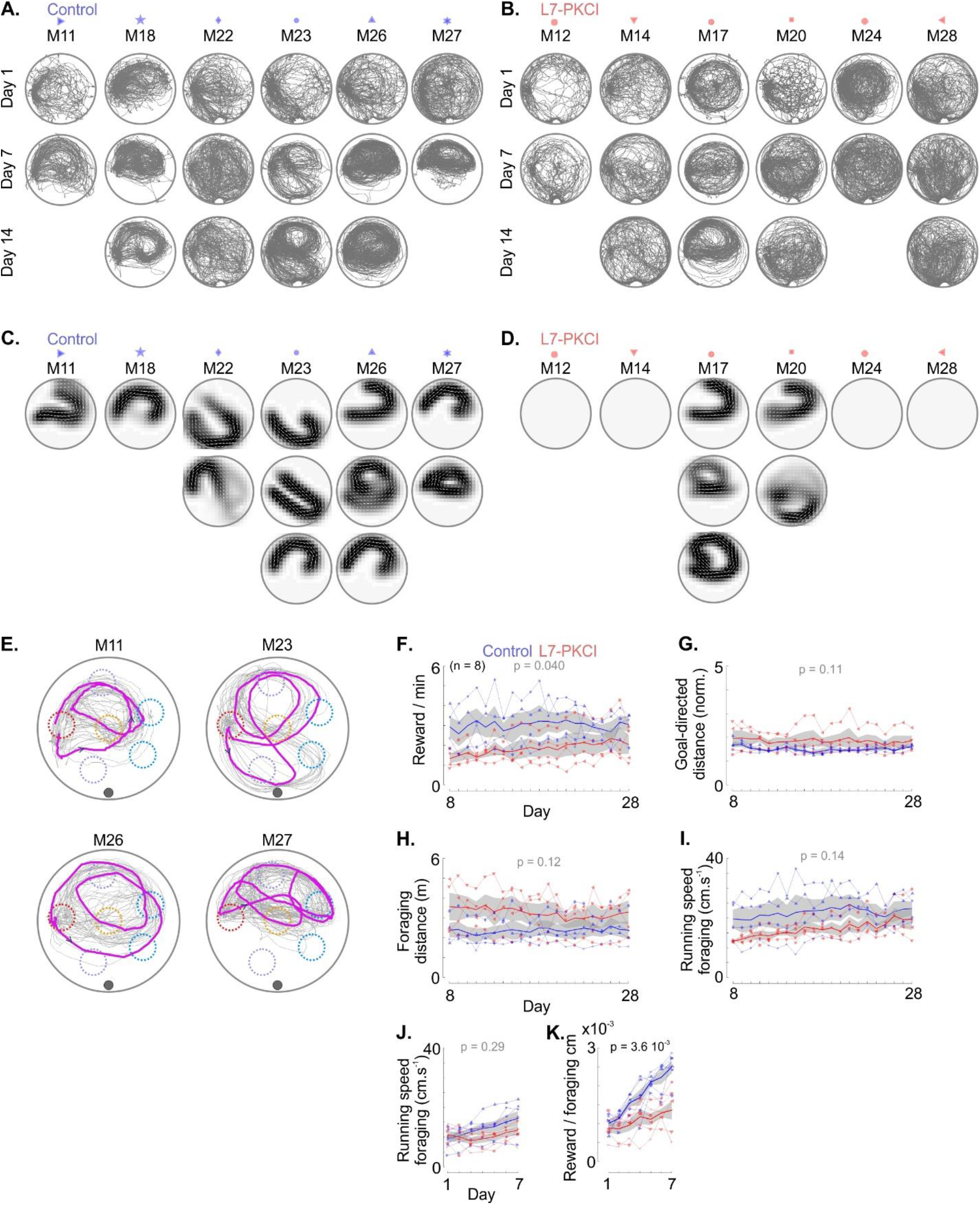
Sequence-based foraging and prolonged training (related to Figure 1). **A.** Examples of trials performed by control mice on the 1^st^ (*top*), 7^th^ (*middle*) and 14^th^ (*bottom*) day of training. Symbols indicate mouse identity throughout the figures. **B**. Same as A for L7-PKCI mice. **C.** Templates of spatial sequence identified by hierarchical clustering in all control mice. Columns correspond to different animals; lines correspond to multiple templates found in the same animal. Symbols indicate mouse identity throughout the figures. **D**. Same as in C for L7-PKCI mice. Blank matrices indicate mice for which no spatial sequence was identified. Arrows indicate the direction of motion. **E.** Example of single-trial trajectories (*purple*) where mice repeated only part of the spatial sequence when they did not obtain the foraging reward. **F.** Number of rewards collected across later days of training (*red*: L7-PKCI, n = 4; *blue*: control, n = 4, mixed GLM, LR test of genotype effect: Δχ^2^(1) = 4.22, p = 0.04). *Thick curve*: median +/− s.e.m.. **G.** Same as in F for the median distance traveled during the goal-directed phase (mixed GLM, LR test of genotype effect: Δχ^2^(1) = 2.52, p = 0.11). The goal-directed distance was normalized in each trial by the shortest distance between the position where the animal received the foraging reward and the position where it first entered the goal zone. **H.** Same as in F for the median distance traveled during the foraging phase (mixed GLM, LR test of genotype effect: Δχ^2^(1) = 2.44, p = 0.12). **I.** Same as in F for the median of running speeds during the foraging phase (mixed GLM, LR test of genotype effect: Δχ^2^(1) = 2.17, p = 0.14). **J.** Median of running speeds during the foraging phase across the first 7 days of training for L7-PKCI (*red*) and control (*blue*) mice (mixed GLM, LR test of genotype effect: Δχ^2^(1) = 1.25, p = 0.29). *Thick curve*: median +/− s.e.m.. **K.** Number of reward per minute normalized by the running speed during foraging. This ratio is thus equivalent to a measure of performance per distance unit, which is independent of running speed. L7-PKCI mice still exhibit a lower performance than control mice (mixed GLM, LR test of genotype effect: Δχ^2^(1) = 8.45, p = 3.6 10^−3^), demonstrating that differences in running speed cannot explain the lower performance of L7-PKCI mice.

**Figure S2.**
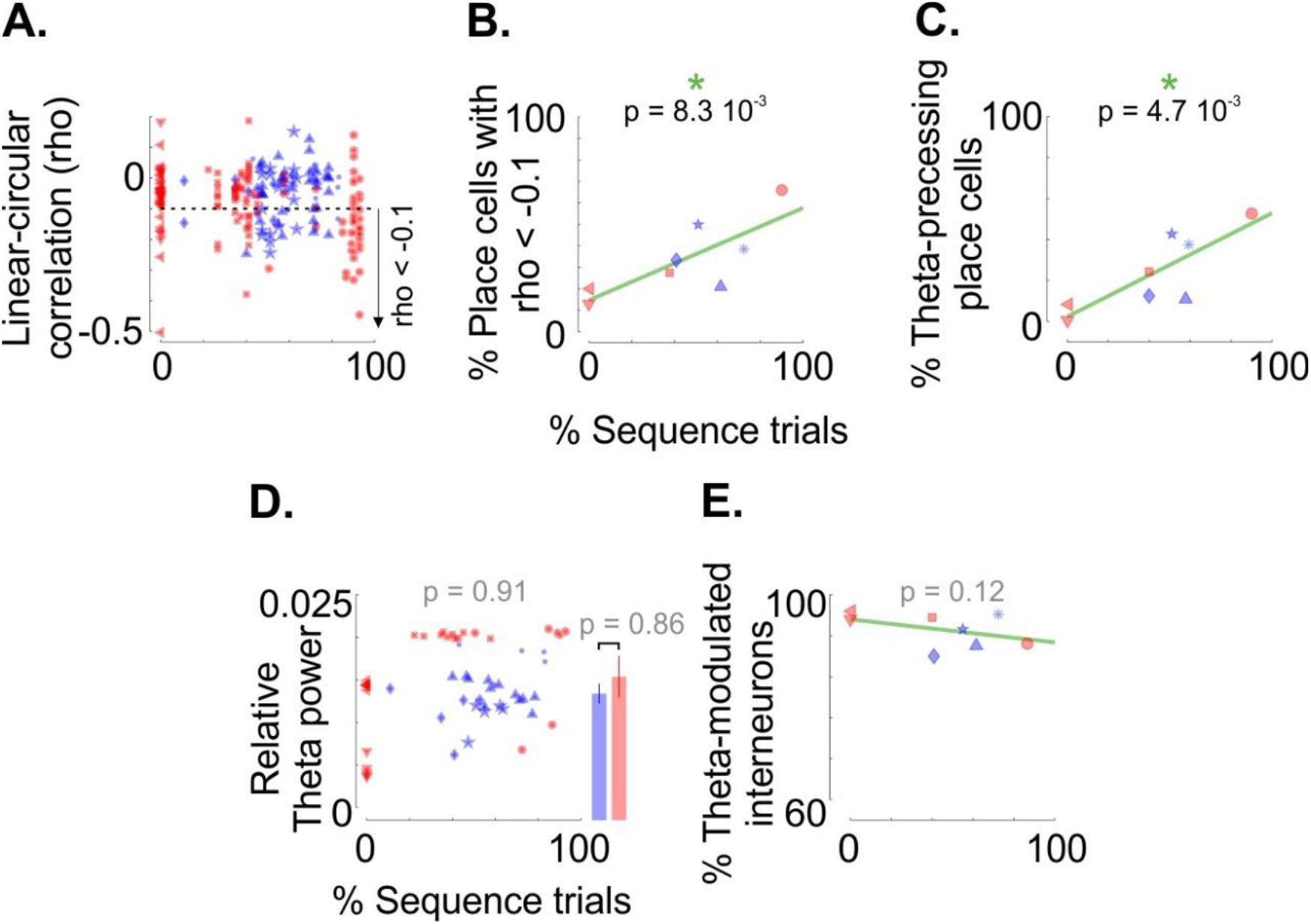
Theta coupling and percentage of sequence trials (related to Figure 2). **A.** Linear-circular correlation (rho) as a function of the percentage of sequence trials for all place cells. Dotted line: −0.1 threshold used in B. **B.** The percentage of place cells with a linear-circular correlation coefficient smaller than −0.1 is significantly correlated with the percentage of sequence trials (mixed GLM, LR test for the fraction of sequence trial: Δχ2(1) = 6.96, p = 8.7 10-3). **C.** Percentage of place cells that exhibited significant theta-phase precession, after selecting only place field traversals that were fast, straight and going through the center of the place field (mixed GLM, LR test for the fraction of sequence trial: Δχ2(1) = 7.99, p = 4.7 10-3). **D.** Theta power as a function of the percentage of sequence trials for L7-PKCI (*red*) and control (*blue*) mice across recording days (mixed GLM, LR test for the fraction of sequence trial: Δχ^2^(1) = 0.014 p = 0.91). Theta power was measured as the mean power between 6 and 9 Hz, normalized by the sum of the power spectrum across all frequencies up to 40Hz. **E.** Fraction of interneurons modulated by theta phases as a function of the percentage of sequence trials for L7-PKCI (*red*) and control (*blue*) mice (mixed GLM, LR test for the fraction of sequence trial: Δχ^2^(1) = 2.48, p = 0.12).

**Figure S3.**
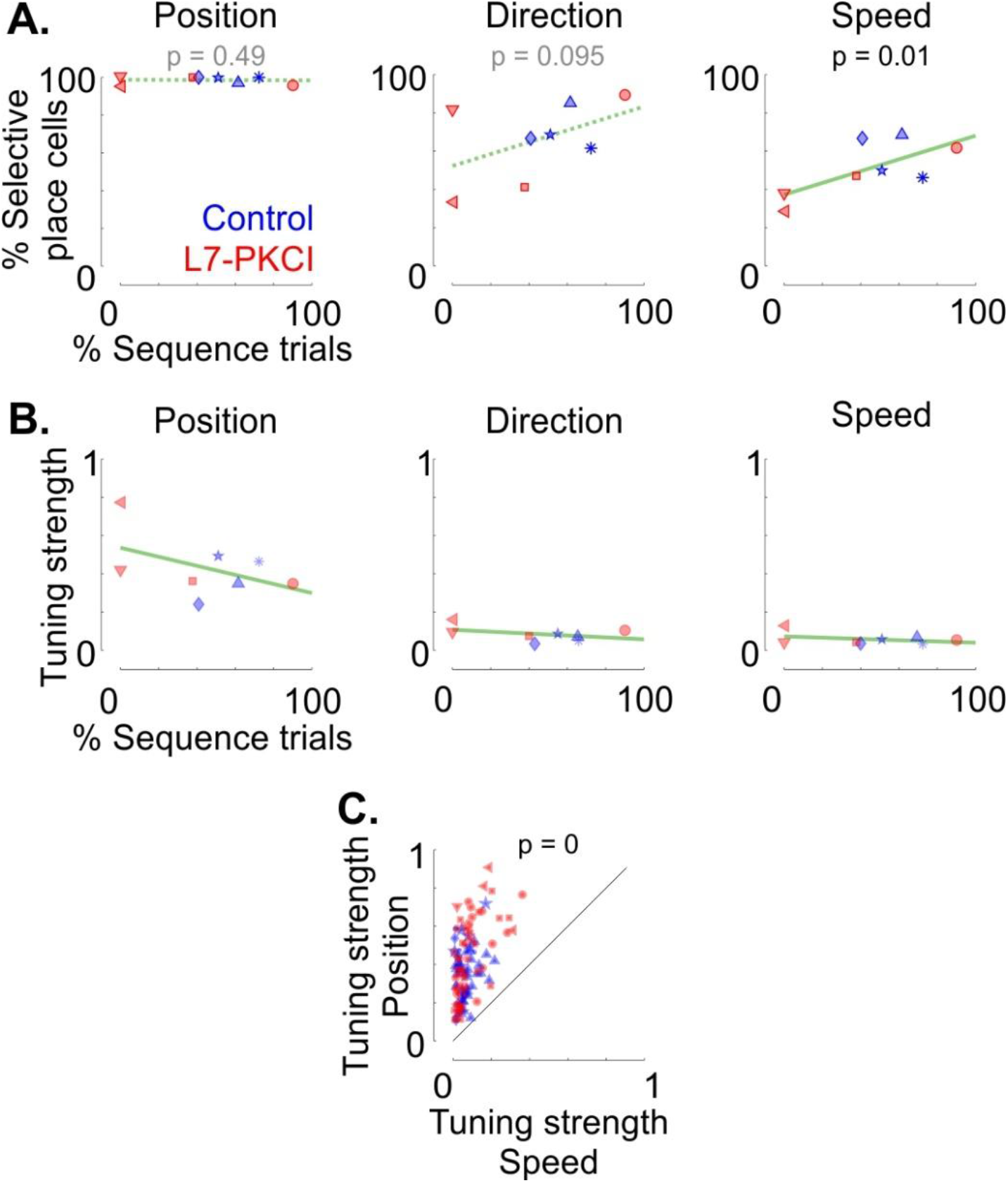
Sequence-based foraging behavior and tuning to positions, direction and speed (related to Figure 3). **A**. Fractions of place cells that were identified as selective to spatial positions (*left*), movement direction (*middle*) or speed (*right*) in L7-PKCI (*red*) and control (*blue*) mice, as a function of the percentage of sequence trials. P-values indicate the significance of the effect related to the fraction of sequence trials (mixed GLM, LR test). **B.** Median across place cells of the strength of tuning to positions (*left*), direction (*middle*) and speed (*right*) for L7-PKCI (*red*) and control (*blue*) animals, as a function of the percentage of sequence trials. **C.** Tuning strength to position as a function of the tuning strength to running speed. The p-value indicates the significance of the Position vs Speed effect (mixed GLM, LR test).

**Figure S4.**
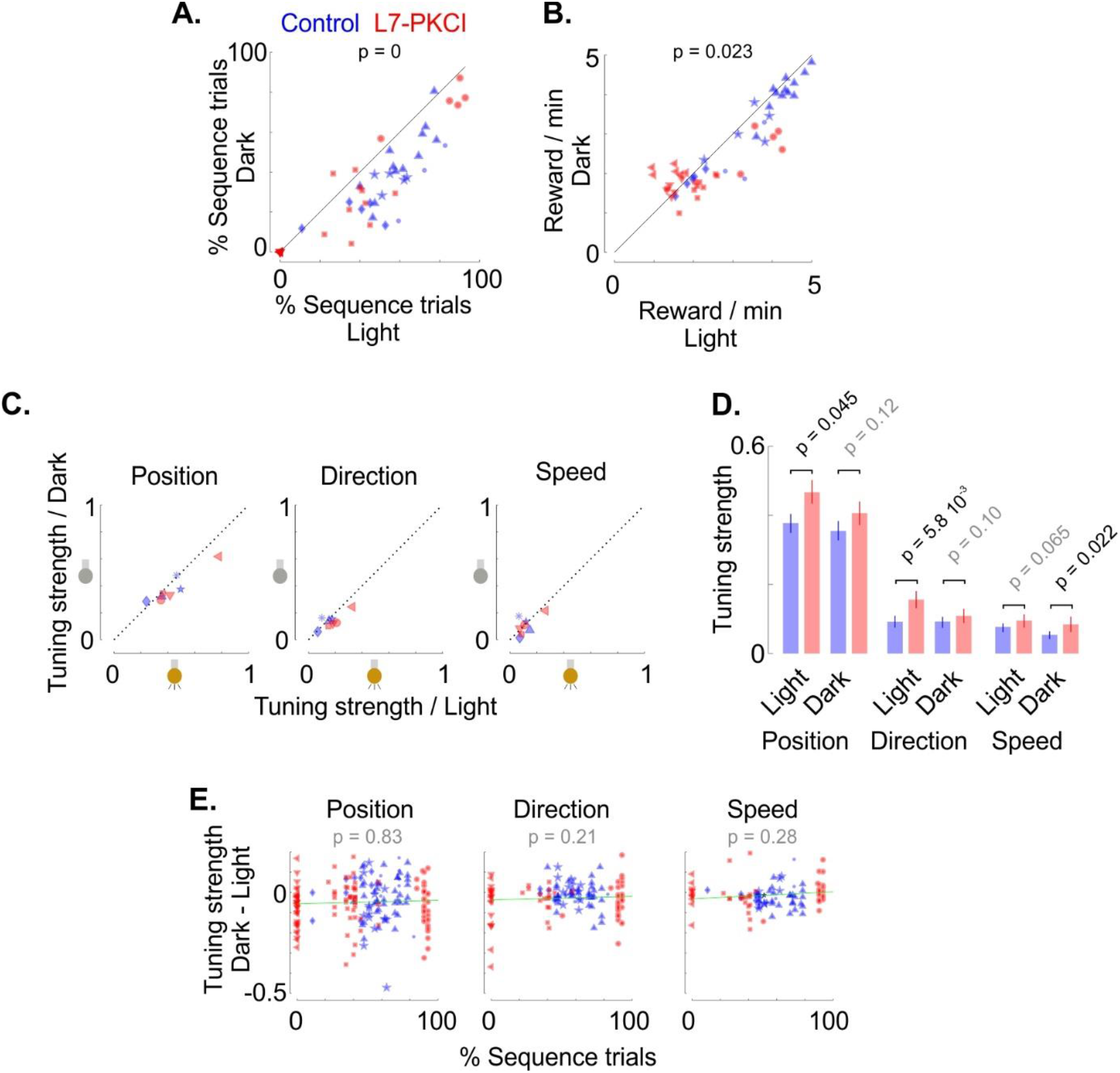
Comparison of performances and place cells tuning between light and dark conditions (related to Figure 4). **A.** Comparison of the fraction of sequence trials between dark and light conditions (mixed GLM, LR test for the effect of light/dark condition: Δχ^2^(1) = 210.56, p = 0; LR test for the interaction between light/dark condition and genotype: Δχ^2^(1) = 1.31, p = 0.25). **B.** Comparison of the number of collected rewards between dark and light conditions (mixed GLM, LR test for the effect of light/dark condition: Δχ^2^(1) = 5.10, p = 0.023; LR test for the interaction between light/dark condition and genotype: Δχ^2^(1) = 0.23, p = 0.63). **C.** Comparison of the median across place cells of tuning to position (*left*), movement direction (*middle*) or speed (*right*) between light and dark conditions, for L7-PKCI (*red*) and control (*blue*) mice. **D.** Tuning strength of L7-PKCI and control place cells for position, direction and speed during light and dark conditions. P-values indicate the significance for the genotype effect (mixed GLM, LR test). **E.** Difference between the tuning strength measured in the dark and in the light (Dark - Light) as a function of the percentage of sequence trials for Position, Direction and Speed tuning. P-values indicate the significance for the interaction between the effect of the light/dark condition and the percentage of sequence trials (mixed GLM, LR test).

